# Metabolic and metagenomic profiling of hydrocarbon-degrading microorganisms obtained from the deep biosphere of the Gulf of México

**DOI:** 10.1101/606806

**Authors:** Aldo Moreno-Ulloa, Victoria Sicairos Diaz, Javier A. Tejeda-Mora, Marla I. Macias Contreras, Fernando Díaz Castillo, Abraham Guerrero, Ricardo Gonzales Sanchez, Rafael Vazquez Duhalt, Alexei Licea-Navarro

## Abstract

Marine microbes are capable of degrading hydrocarbons; however, those inhabiting the deep biosphere (>1000 m) remain largely unexplored. Microbial metabolism could lead to the generation of diverse chemistries (some with therapeutic activities), but the impact of using hydrocarbons as the sole source of microbial energy in the synthesis of metabolites, remains obscure. Here, we investigated the metagenomic and metabolomic profile of two deep-marine sediments (>1 200 m deep, designated as A7 and B18) collected from the Gulf of México (GM) when grown for 28 days with a simple mixture of 4 hydrocarbons and complex hydrocarbon mixture (petroleum API 40) as their sole source of energy. State of the art techniques and analysis (e.g., Global Natural Products Social Molecular Networking, network annotation propagation [NAP], and MS2LDA) were used to describe the chemistries associated to the microbial utilization of hydrocarbons. The metagenomic sequencing analysis suggests a predominant abundance of *Proteobacteria* in environmental and API 40-enriched samples, while the abundance of *Pseudomonas* increased after microbial growth with API 40. The metabolomic analysis suggests the presence of diverse chemistries predominantly associated with lipid and lipid-like and phenyl propanoids and polyketides superclass (Classyfire annotation). Hydrocarbon derivatives were detected as carboxylic acids (e.g., azelaic and sebacic acid) or alcohols, while non-hydrocarbon related chemistries were also detected including tetracycline-related metabolites and sphinganines. Our study provides valuable chemical and microbiological information of microbes inhabiting one of the most understudied ecosystems in the earth, the deep marine biosphere.

## 1. Introduction

The increasing number of oil platforms in the Gulf of México (GM) represents an environmental risk. As a consequence, the GM it has been the scenario of the world’s largest accidental release of oil into the ocean (i.e., Deepwater Horizon oil spill) with approximately 4.9 million barrels of crude oil flowed into marine sediments (Team 2010). It is well known that marine microbes possess the capacity to metabolize hydrocarbons and thereby contribute to the bioremediation of oil spills into the ocean (Kostka et al 2011). However, most studies have focused on the ocean surface and water column environments (Brooijmans et al 2009). By comparison, deep-marine (>1000 m) sediments remain poorly studied pertaining to the metabolic capacities of microbes towards hydrocarbons. Deep-sea sediments harbor diverse and abundant microbial communities comparable to those in seawater (Kallmeyer et al 2012). Studies have shown the enrichment of selected bacteria and genes associated with aliphatic and aromatic metabolism in sediments after 5-6 months of the Deepwater Horizon oil spill (Mason et al 2014).

The limited research on hydrocarbon degradation or metabolism of microbes in deep-marine sediments has been inferred, primarily, from bacterial communities or metagenomes found at those environments (Brooijmans et al 2009, Mason et al 2014). Few studies have focused on hydrocarbon-related degradation products by microbial metabolism using selected types of hydrocarbons (e.g., aliphatic and aromatics) by employing gas chromatography coupled to mass spectrometry (MS) and specific hydrocarbon standards (Bacosa et al 2018, Kimes et al 2013). Despite the abundant information on individual microorganisms (e.g., bacteria) or bacterial consortiums capable of degrading hydrocarbon compounds, scarce information is available about the metabolism of microbes inhabiting the deep-sea biosphere. This could be attributed, in part, due to the complexity of microbial metabolism, technical limitations to globally assess the chemistries linked to microbes, and access to deep-sea environments. Noteworthy, reports have described the capability of marine microbes to synthesize novel molecules, distinct to those of their terrestrial counterpart, with prominent therapeutic activities (Blunt et al 2018). It is estimated that over twenty thousand molecules have been discovered from marine sources (Gerwick and Moore 2012)

High-resolution tandem MS (HR-MS/MS)-based metabolomics has demonstrated to be a powerful tool to inventory the chemistry derived from microbe’s cultures (Floros et al 2016). This technique could generate a large amount of data containing known and new molecules that when coupled to appropriate computational tools could lead to a rapid dereplication (i.e., identification of known molecules) process (Mohimani et al 2018). In this regard, the Global Natural Product Social (GNPS) molecular networking platform has contributed to the abovementioned process as well as to the discovery of new chemical entities in a more rapid and precise manner compared to other conventional computational approaches (Quinn et al 2017). This platform analyzes the MS/MS or MS2 spectral data, and based on fragmentation pattern similarity generates molecular networks whereby compound dereplication, through spectral library matching, enriches the networks allowing to infer molecular families. Molecular networking has been successfully implemented to globally visualize the metabolome derived from individual and grouped bacterial strains (Floros et al 2016), study small molecules within a specific pathway (Vizcaino et al 2014), and for targeted-isolation of new chemical entities (Kang et al 2018), among others.

To our knowledge, our work is the first study on the global metabolome of marine microbes when grown with hydrocarbons as their sole source of carbon. Here, we applied molecular networking and complementary ‘‘state of the art’’ *in-silico* dereplication strategies to two selected enriched hydrocarbon-degrading microbes from deep-marine sediments (>1,200 m) of the GM. This study provides valuable chemical and microbiological information of one of the most understudied earth ecosystem, the deep-marine biosphere.

## Materials and methods

### Sample collection

Two superficial sediment samples (0-10 cm) from water depths of 1 265 m (designated as B18, 23°54′57.6″N 86°47′28.8″W) and 3 500 m (designated as A7, 24°57′37.74″N 90°0′52.5″W) were collected, using a Reineck box core (50 × 50 cm), in August 2015 during the GM oceanographic campaign XIXIMI-4 (Figure 2). Samples were collected and stored in sterile cap tubes of 10 cm^3^ at 0-4 °C while on board during the ocean campaign, and then transported to the in-land laboratory for storage at −80 °C until processing.

**Figure 1.**
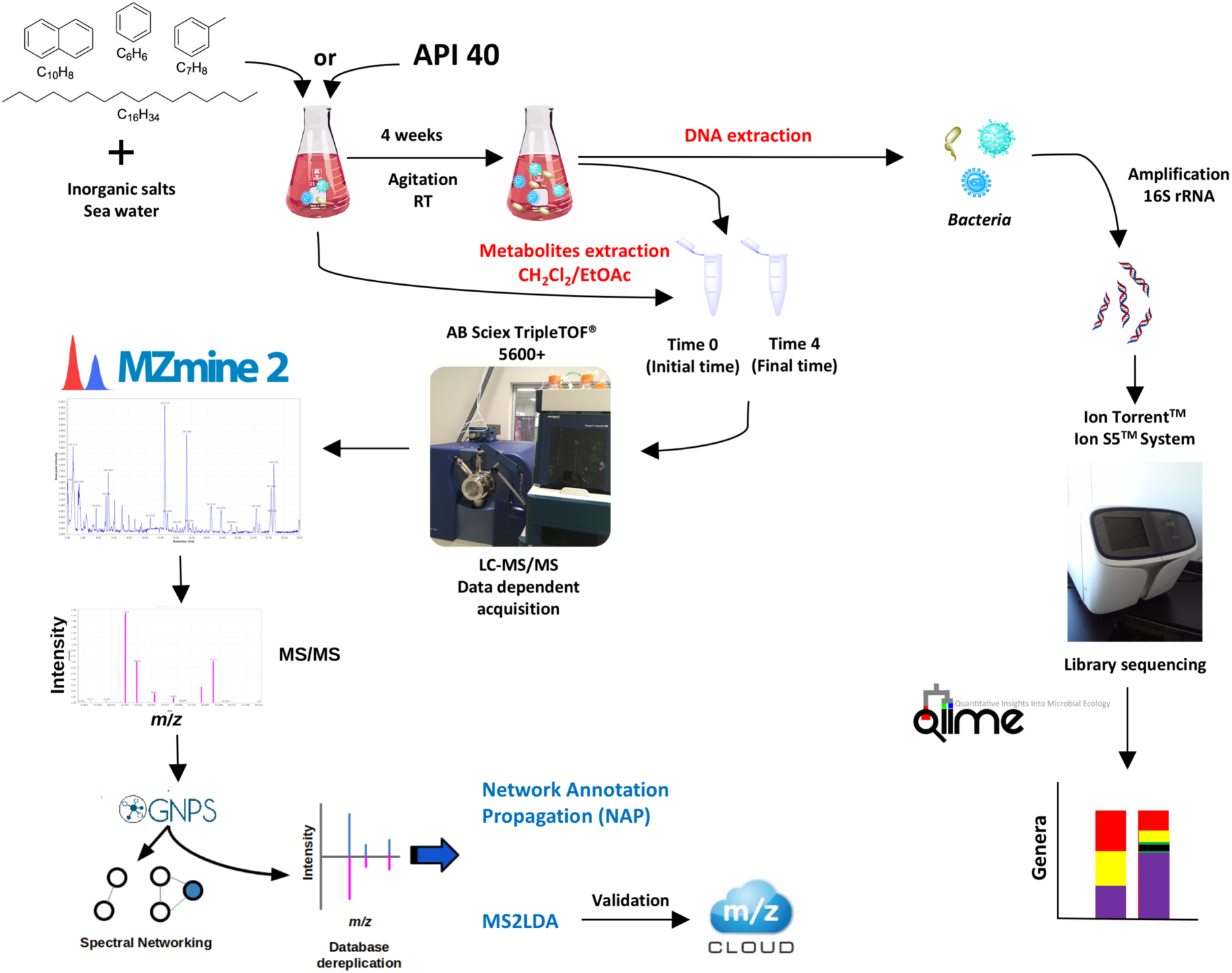
Illustration of the methodology followed in this study.

**Figure 2.**
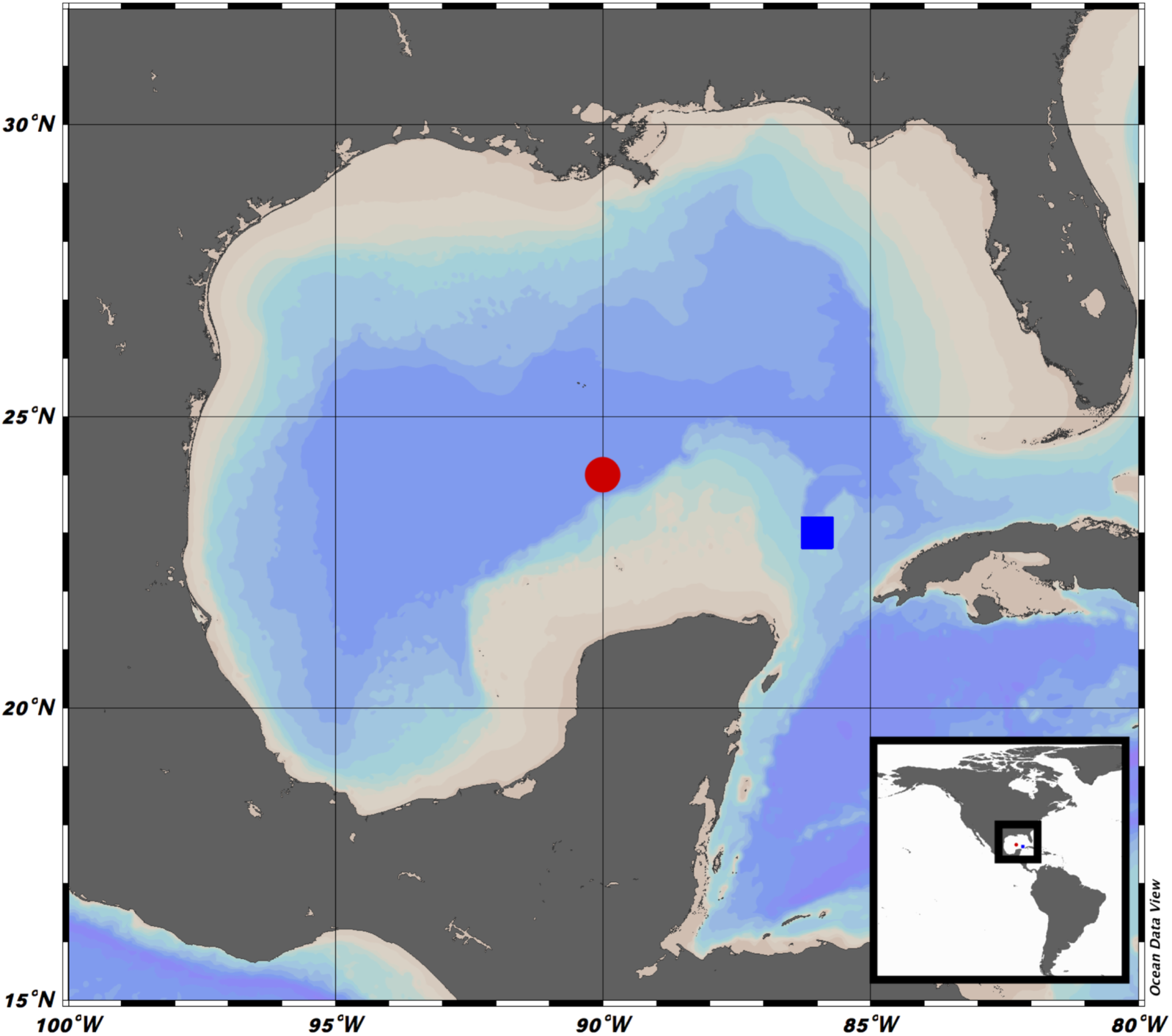
Sampling locations in the Gulf of México. Two superficial sediment samples (0-10 cm) from water depths of 1 265 m (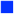, designated as B18, 23°54 ′ 57.6 ″ N 86°47 ′ 28.8 ″ W) and 3 500 m (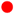, designated as A7, 24°57’37.74"N 90°0’52.5"W).

### Marine sediments culture conditions

Marine sediments were cultured with hydrocarbons as the sole source of carbon to enrich the culture with hydrocarbon-degrading microbes as following: samples were cultured in a mineral medium containing inorganic salts (4 g L^−1^ NaNO_3_, 2 g L^−1^ K_2_HPO_4_, 1 g L^−1^ MgSO_4_.7H_2_O, 1 g L^−1^ KCl, 0.02 g L^−1^ FeSO_4_.7H_2_O), trace elements (0.26 g L^−1^ H_3_BO_3_, 0.50 g L^−1^ CuSO_4._5H_2_O, 0.50 g L^−1^ MnSO_4_.H_2_O, 0.06 g L^−1^ MoNa_2_O_4_.H_2_O, 0.70 g L^−1^ ZnSO_4_.7H_2_O, 0.0185 g L^−1^ CoCl_2_, pH 7±0.2) (Abalos et al 2004) dissolved in three parts of sea water (filtered by a 0.2 µm membrane) and one part of distilled water supplemented with crude oil (1% v/v) as the sole source of carbon. The cultures were maintained, under darkness for four weeks at 19±1 °C. The microbial growth was estimated as the most-probable-number method (MPN) (Coulon et al 2005). After this enrichment step, hydrocarbon-degrading microbes were grown in OxiTop^®^ flasks for another four weeks with two different carbon sources; (I) crude oil (API 40) and (II) synthetic oil (SO, in mole basis, toluene 72.15%, benzene 8.27%, naphthalene 11.33%, and hexadecane 8.25%). For I, flasks contained mineral medium (43.226 mL), API 40 oil (0.4% v/v), and microbial-enriched broth inoculum (100 μL). For II, flasks contained mineral medium (43.313 mL), 87 μL of SO, and microbial-enriched broth inoculum (100 μL). The total volume in both cases was 43.5 mL. Different control cultures were performed: (III) flasks containing mineral medium and either API 40 oil or SO without microbial-enriched broth; (IV) flasks containing mineral medium and microbial-enriched broth without API 40 or SO. After the OxiTop® flasks were filled, a carbon-dioxide trap was fixed in each flask containing three pellets of KOH. Bottles were then capped with OxiTop^®^ heads and placed under magnetic stirring to ensure homogeneity at 19±1 °C for four weeks. All experiments were performed in triplicate. Data collection was started immediately.

### Sequencing and Bioinformatic Analysis

DNA was extracted from environmental and final time samples. The total DNA was extracted by a PowerMarx® Soil (MoBio, Carlsbad, CA, USA) DNA isolation kit following the manufacturer’s instructions. The concentration and purity of the extracted DNA were determined using a NanoDrop™ Lite spectrophotometer (Thermo Scientific, USA).

### Metagenomic Sequencing and Analyses

16S rRNA sequencing was focused in 7 hypervariable regions (V2-V4 and V6-V9) generated from 1 μg of purified DNA, implemented in PGM (ion Torrent technology). The sequencing was carried out according to the Ion 16S ™ Metagenomics Kit (A26216; manual: MAN0010799, TermoFisher).

Sequences with 100 to 400 bp long, and ≥25 of quality (QC) were selected for the analysis. The filtered reads were used to detect chimeric reads by VSEARCH (Rognes et al 2016) using the Gold database (http://drive5.com/uchime/uchime_download.html). The operational taxonomic units (OTUs) were generated by the UCLUST method, using the close reference model implemented at 97% of similarity with Silva database release 123 (Quast et al 2013). The alpha diversity metrics were implemented using the Shannon index; differences among microbial communities were determined using UniFrac and Jackknifed UPMGA. All analyses were implemented in QIIME version 1.9 (Caporaso et al 2010).

### Metabolites extraction

Initial and final time samples were extracted by liquid-liquid extraction with dichloromethane and ethyl acetate (50:50). Then, the solvent extracts were dried over Na_2_SO_4_, concentrated in a vacuum rotary evaporator, and finally dried under a gentle nitrogen stream. Samples were stored at −20 °C until processing.

### Liquid chromatography coupled to HR-MS/MS (LC-HR-MS/MS)

Dried extracts were re-dissolved in 5% acetonitrile (ACN) in water with 0.1% formic acid and analyzed using an Eksigent nanoLC^®^ 400 system (AB Sciex, Foster City, CA, USA) with a HALO Phenyl-Hexyl column (0.5 × 50 mm, 2.7 μm, 90 Å pore size, Eksigent AB Sciex, Foster City, CA, USA). The separation of metabolites was performed using gradient elution with 0.1% formic acid in water (A) and 0.1% formic acid in ACN (B) as mobile phases at a constant flow rate of 5 μL/min. The gradient started with 5% B for 1 min, changing to 100% B in 26 min and held constant over the next 4 min. Then, solvent composition was changed back to 5% B over 0.1 min. Four minutes post run with 100 % mobile phase A were applied to ensure column re-equilibration. One blank was run between sample injections to minimize potential carryover. The eluate from the LC was delivered directly to the TurboV source of a TripleTOF 5600+ mass spectrometer (AB Sciex, Foster City, CA, USA) using electrospray ionization (ESI) under positive ion. ESI source conditions were set as following: IonSpray Voltage Floating, −5500 V; Source temperature, 350 °C; Curtain gas, 20 arbitrary units; Ion Source Gas 1, 40 arbitrary units, Ion Source Gas 2, 45 arbitrary units and; Declustering potential, −100. Data was acquired using information-dependent acquisition (IDA) with high sensitivity mode selected, automatically switching between full-scan MS and MS/MS. The accumulation time for TOF MS was 0.25 s/spectra over the *m/z* range 100-900 Da and for MS/MS scan was 0.05 s/spectra over the *m/z* 50-900 Da. The IDA settings were: charge state +1 to +2, intensity 125 cps, exclude isotopes within 6 Da, mass tolerance 50 mDa, and a maximum number of candidate ions 20. Under IDA settings, the ‘‘exclude former target ions’’ was set as 15 s after two occurrences and ‘‘dynamic background subtract’’ was selected. The collision energies were varied to optimize sensitivity. The instrument was automatically calibrated by the batch mode using appropriate positive TOF MS and MS/MS calibration solutions before sample injection and after injection of two samples (<3.5 working hours) to ensure a mass accuracy of <5 ppm for both MS and MS/MS data.

### LC-HR-MS/MS data processing and analysis

Raw files (.wiff and wiff.scan) were converted to .mzML using ProteoWizard (Holman et al 2014) and processed with MZmine 2.34 (Pluskal et al 2010). Centroided mass detection was done keeping the noise level at 1 and 0.01 for MS1 and MS2, respectively. Chromatograms were built using the ADAP algorithm (Myers et al 2017) by inputting the following parameters: intensity threshold of 5.0, minimum highest intensity of 5.0 and m/z tolerance of 10 ppm. Chromatogram deconvolution was done using ADAP algorithm, and chromatograms were deisotoped. Peaks initial times were removed from final times samples by using the peaks row filter option after peak alignment (join algorithm). Finally, peaks with MS/MS data were exported (.MGF file for GNPS) for further analysis.

MGF files were analyzed by the Global Natural Products Social Molecular Networking (GNPS) Data Analysis workflow (https://gnps.ucsd.edu) to create a molecular network (Wang et al 2016). The data were filtered by removing all MS/MS peaks within +/-17 Da of the precursor m/z. MS/MS spectra were window filtered by choosing only the top six peaks in the +/- 50Da window throughout the spectrum. The data was then clustered with MS-Cluster with a parent mass tolerance of 0.02 Da and a MS/MS fragment ion tolerance of 0.02 Da to create consensus spectra. A network was then created where edges were filtered to have a cosine score above 0.6 and more than 5 matched peaks. Further edges between two nodes were kept in the network if and only if each of the nodes appeared in each other’s respective top 10 most similar nodes. The spectra in the network were then searched against GNPS’ spectral libraries. The library spectra were filtered in the same manner as the input data. All matches kept between network spectra and library spectra were required to have a score above 0.7 and at least 5 matched peaks. Molecular networking data was visualized using Cytoscape 3.6.1 (Shannon et al 2003). The generated molecular network was further analyzed using the Network Annotation Propagation (NAP) tool to annotate, by *in silico* predictions, candidates for individual spectra and predict the chemical class of molecular networks or node clusters (based on the most predominant chemical class within a network) (da Silva et al 2018). The parameters used for NAP were as follows: ten first candidates for consensus score, positive acquisition mode, exact mass searches within 10 ppm and GNPS and HMDB databases.

Furthermore, the clustered .mgf file generated by GNPS was subjected to MS2LDA (https://MS2LDA.org) (van der Hooft et al 2016) for extracting Mass2motifs (M2M). MS2LDA tool allows to discover (in an unsupervised manner) groups of neutral losses and mass fragments termed M2M linked to substructures within the datasets. The parameters used were set as follows: input format .mgf, *m/z* tolerance 5.00 pm, minimum MS2 intensity 0.1 a.u., number of iterations 1000, and number of M2M 80.

## Data availability

### GNPS molecular network

https://gnps.ucsd.edu/ProteoSAFe/status.jsp?task=133fac1cd5ca46c28caea1362babc556

### NAP molecular networks

All submitted cluster indexes are in an excel table in the supplementary information

### MS2LDA network

http://ms2lda.org/basicviz/summary/899/; A7 and B18 combined data

## Results

### Growth of microorganisms of deep-marine sediments with hydrocarbons as their sole source of energy

**Figure 3A** shows the microbial growth of A7 and B18 with API 40 as per the oxygen consumption rate analysis. There were significant differences in the oxygen consumption rate between A7 and B18, for instance, A7 and B18 show a lag time of approximately ∼4 days and ∼6 days, respectively. Visually, it appears that A7 follows a two-step in the oxygen consumption pattern with two exponential phases from ∼6 to 10 days (63.72±4.28 mg/mL•day^−1^) and ∼10 to 28 days (18.53±0.36 mg/mL•day^−1^) that were fitted with a linear regression (r^2^>0.59 and r^2^>0.79). On the other hand, B18 show a lag time of ∼6 days and a unique exponential phase from 10 to 28 days (33.38±0.65 mg/mL•day^−1^) also fitted with a linear regression (r^2^>0.79). The first slope of A7 indicates a significant higher oxygen consumption rate than B18 (63.72±4.28 mg/mL•day^−1^vs. 33.38±0.65 mg/mL•day^−1^, p<0.0001) (**Figure 3B**), whereas, after 10 days, A7 shows a lower growth rate compared to B18 (18.53±0.36 mg/mL•day^−1^ vs. 33.38±0.65 mg/mL•day^−1^, p<0.0001) (**Figure 3C**).

**Figure 3.**
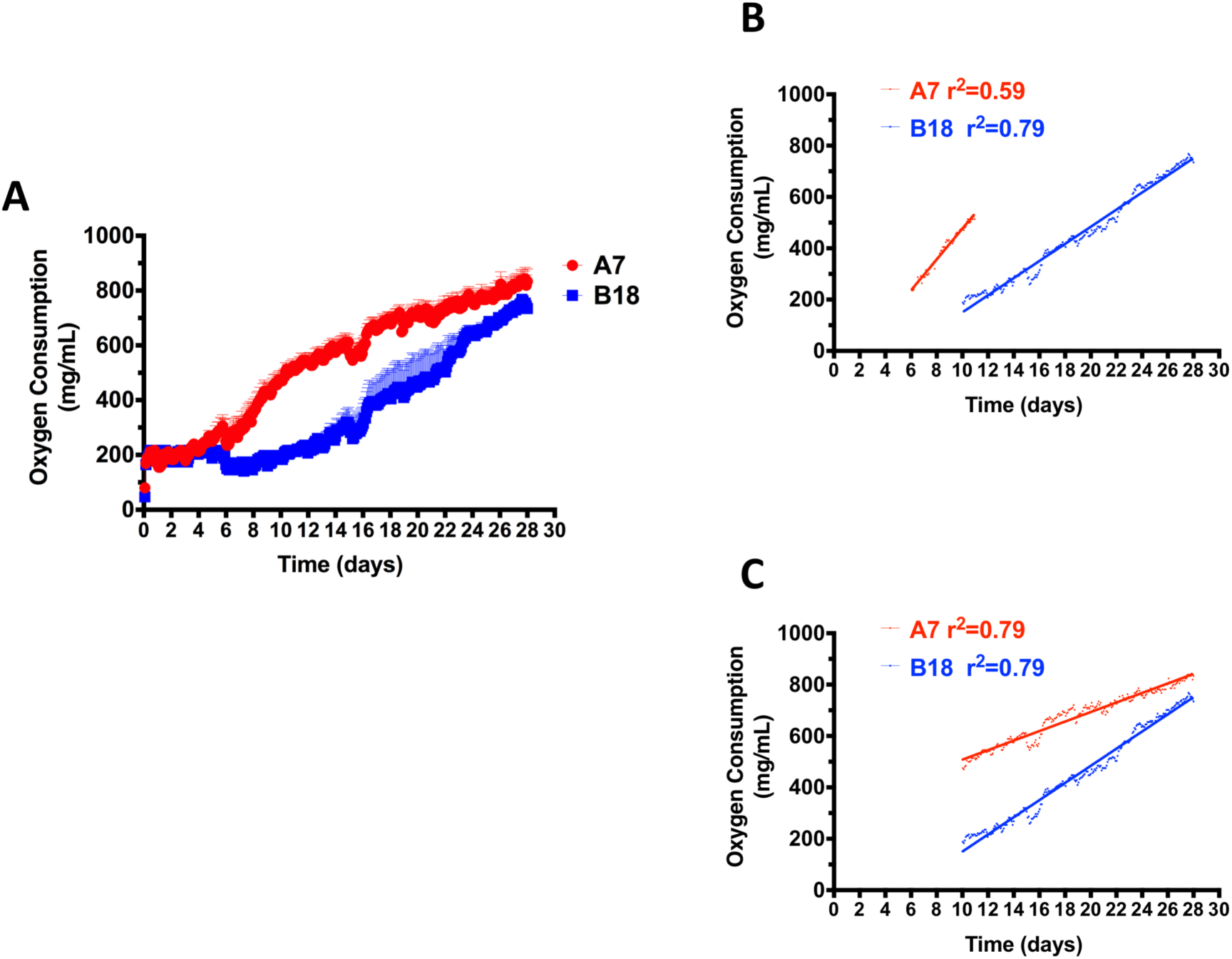
Growth standardization of microorganisms from deep-marine sediments. (A) Microbial growth of A7 and B18 was evaluated in media containing 4 mg/L of sodium nitrate and 1% v/v API 40. Linear oxygen consumption within the growth curve were fitted with a linear regression after a lag time of 0-6 days. (B) Linear regression analysis of the first exponential growth phase (6 to 10 days) of A7 compared to that of B18 (10 to 28 days). (C) Linear regression analysis of the second exponential phase of A7 (10 to 28 days) compared to that of B18 (10 to 28 days). Detail calculations are explained within the results section. All experiments were performed in triplicate.

### Microbial communities of environmental deep-marine sediments and API 40-enriched sediment samples

The bioinformatics analysis led to the detection of 1 934 operational taxonomic units (OTUs) associated with bacteria, whereby 282 were classified up to genera. Globally, most of the taxon detected among environmental and API 40-enriched samples were associated to Proteobacteria (64.8%), Firmicutes (9.6%), Acidobacteria (7.3%), Actinobacteria (6.9%), and others. Of those 282 genera detected, 28 genera were shared among environmental and API 40-enriched samples. Environmental samples presented with 182 genera and of those, a 60.6% were characteristic of this group when compared to API 40-enriched samples. On the other hand, API 40-enriched samples presented with 100 genera and of those a 28.3% were characteristics of this group when compared to their counterpart. Analysis at the genus level, shows that the most abundant genera were *Pseudomonas* (16.7%), *Staphylococcus* (4.8%), *Altererythrobacter* (4.5%), *Erythrobacter* (4.2%), *Erythrobacter* (4.2%), *Novosphingobium* (1.4%), *Bacillus* (2%), *Pantoea* (1.8), *Nitrospira* (1.3%), *Calditerrivibrio* (1%), *Propionibacterium* (0.8%), while the remaining genera were present with lower values (**Figure 4**).

**Figure 4.**
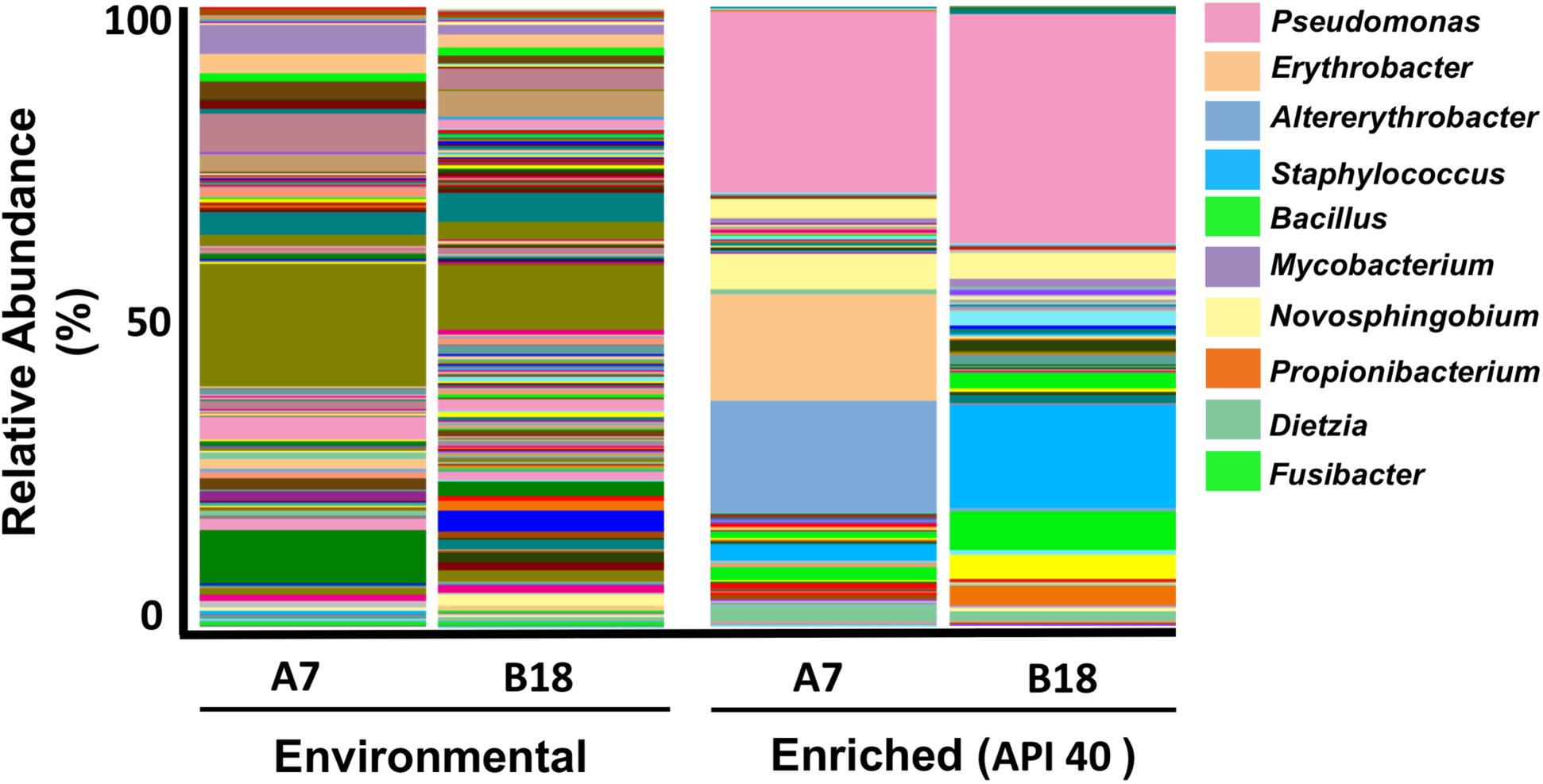
Bacterial community diversity at genus level in environmental deep-marine sediments and incubated with petroleum API 40.

Particularly, 44 (24.2%) genera were shared between A7 and B18 environmental samples, while 75 (41.2%) and 63 (34.6%) were exclusively detected in A7 and B18, respectively. The most abundant phyla in A7 were Proteobacteria (61.9%), Actinobacteria (10.9), Acidobacteria (7.1%), Gemmatimonadetes (6.1%), and others. Analysis at the genus level in A7 suggest the presence of *Nitrospira* (3.6%), *Caldithrix* (1.7%), *Aquibacter* (0.25%), *Nitrospina* (0.26), *Amaricoccus* (0.11%), *Pelagibius* (0.10%), *Nitrosomonas* (0.9%), *Alteromonas* (0.087%), *Pseudoalteromonas* (0.57%), OM60(NOR5) clade (0.18), and H16 (0.53%), while the rest presented with less than 0.06% abundance. In B18 environmental samples, the most abundant phyla were Proteobacteria (56.4%), Acidobacteria (21.2%), Actinobacteria (5.2%), Nitrospirales (3.5%), and others. The most abundant genera were *Nitrospira* (1.91%), Pseudomonas (1.12%), *Pelagibius* (1.79), H16 (1%), Urania-1B-19 (0.85%), *Nitrospina* (0.66%), *Pelagibius* (0.79%), *Pseudoalteromonas* (0.59%), *Aeromonas* (0.52%), and the remaining presented with less than 0.4% abundance.

On the other hand, of the 100 genera detected in API 40-enriched samples; 33 (33%) were detected in both A7 and B18 samples, while 24 (24%) and 43 (43%) of those were exclusively detected in A7 and B18, respectively. The most abundant phyla in A7 were Proteobacteria (83%) and Gammaproteobacteria (34%) and analysis at the genus levels suggest the presence of 57 genera, whereby the most abundant genus was Pseudomonas (29%), Altererythrobacter (18.1%) and *Erythrobacter* (17%), *Novosphingobium* (5.6%), *Pantoea* (3.1%), *Dietzia* (2.9%), *Bacillus* (1.9%), *Staphylococcus* (2.6%), the rest of the genera presented with less than 1% abundance (**Figure 4**). In the case of B18, the most abundant phyla were Proteobacteria (57.4%), Firmicutes (29.5%), Actinobacteria (6.2%), and Deferribacteres (3.9%). Analysis at the genus level suggests the presence of 76 genera, whereby the most abundant genus was *Pseudomonas* (36.8%), *Pantoea* (4.2%), *Nannocystis* (2.2%), *Fusibacter* (2.5%), *Staphylococcus* (16.6%), *Bacillus* (6.2%),*Calditerrivibrio* (3.9%), *Propionibacterium* (3%), *Janthinobacterium* (1.8%), *Dietzia* (1.4%), and *Klebsiella* (1.3%), while the rest of the genus presented with less than 1% abundance (**Figure 4**).

To gain further insight into the microbial community in deep-marine sediments, we evaluate differences in abundance of microbes with reported oil-degrading capabilities, referred as to oil-degrading bacteria (ODB) (Yakimov et al 2007). Globally, 28 genera were detected as ODB (**Figure 5**). The common genera detected among environmental and API 40-enriched samples were *Pseudomonas, Mycobacterium, Stenotrophomonas*, and *Candidatus*. The environmental samples presented with 21 genera (2.1% relative abundance on average), and of those 12 (0.65% relative abundance) and 16 (3.6% relative abundance) were detected in A7 and B18, respectively. Seven genera were shared between A7 and B18. As expected, API-40 enriched samples presented with a higher average relative abundance of ODB (51%). In A7, 13 genera (38.9% relative abundance) were detected while 14 genera (64.5%) were detected in B18, 11 genera were shared between both samples.

**Figure 5.**
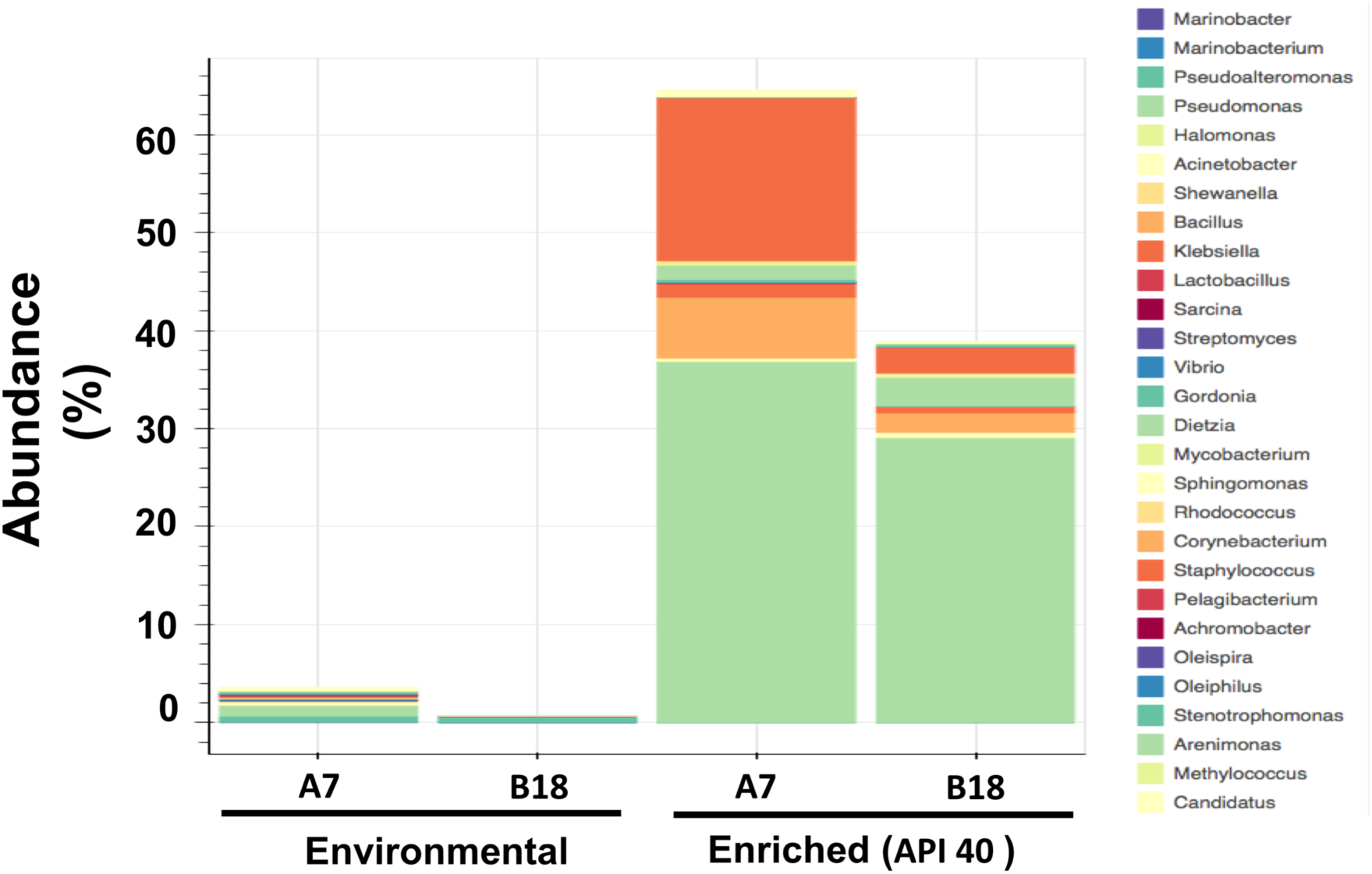
Abundances of genera associated to Oil-Degrading Bacteria (ODB) in environmental deep-marine sediments and incubated with petroleum API 40.

### Metabolites associated with hydrocarbon-degrading microorganisms from deep-marine sediments

All LC-MS/MS data were analyzed by MZmine, whereby for each dataset features detected in initial times were subtracted from final times to retain the features associated with hydrocarbon metabolism. To inventory the chemistry of microorganisms with hydrocarbon degrading capabilities, we submitted LC-MS/MS datasets (processed with MZmine), derived from A7 and B18 grown with both API 40 and SO, to the GNPS web platform (https://gnps.ucsd.edu) (Wang et al 2016). The generated combined molecular network or spectral network comprised 1 501 mass spectral nodes (over a mass range of 113.06-882.687 *m/z*, **Supplementary Figure 1**) organized into 129 independent molecular families (with at least two connected nodes) (**Figure 6A**). Within the global molecular network, the node and border color reflect the type of hydrocarbon (blue for SO and green for API 40) and microbial sample (red for A7 and orange for B18) used, respectively. The ion co-occurrence in two or more conditions is denoted by a red intermittent or star node border, respectively. A 4.8% (n=72) and 0.1% (n=2) of precursor ions co-occur in two and three or more culturing conditions, denoting a high variability in secondary metabolite production between A7 and B18 microbes. The number of unique precursor ions was considerably higher for B18 grown with SO followed by A7 under the same condition. The spectral data was screened against public spectral databases within the GNPS, which led to a 0.4% (n=6) of dereplicated molecules within our molecular network. To further characterize the chemistries of B18 and A7, we employed the NAP tool (da Silva et al 2018) to propagate structural annotations, by *in silico* predictions, throughout the molecular network and automatically classify the networks as per the most predominant chemical superclass using the ClassyFire tool (Djoumbou Feunang et al 2016). Our data show a diverse type of secondary metabolites, particularly lipid and lipid-Like molecules, followed by phenyl propanoids and polyketides (**Figure 6B**). As a complementary approach to NAP, we also employed the MS2LDA computational tool aiming at discovering co-occurring fragments and neutral losses (named M2M) within the MS2 data (using the same .mgf file as in GNPS and NAP), which provides chemical information at the substructure level (van der Hooft et al 2016). However, due to the challenging task of characterizing the entire metabolome of microbes derived from untargeted analysis and the metabolic complexity of potential non-previously described microbes, we applied the NAP/MS2LDA-driven metabolite annotation of selected molecular networks (i.e., enriched-networks with standards or spectral library hits) as described by Kang et al., (Kang et al 2019). Tetracycline standard clustered within a network of 29 nodes (**Figure 7A**), whereby MS2LDA and manual inspection analysis, using fragment search by mzcloud (www.mzcloud.org), led to the putative identification of tetracycline-related chemistries as per the motif M2M_14. The fragment 410.1225 *m/z* suggests a tetracycline-related core in the connected node (444.698 *m/z*) to tetracycline, while the other nodes do not share this fragment but do others (Figure 7A). Likewise, the assigned superclass to ‘‘phenyl propanoids and polyketides’’ of such network by NAP (i.e., through Classyfire) is in agreement to the chemical classification of tetracyclines (Djoumbou Feunang et al 2016).

**Figure 6.**
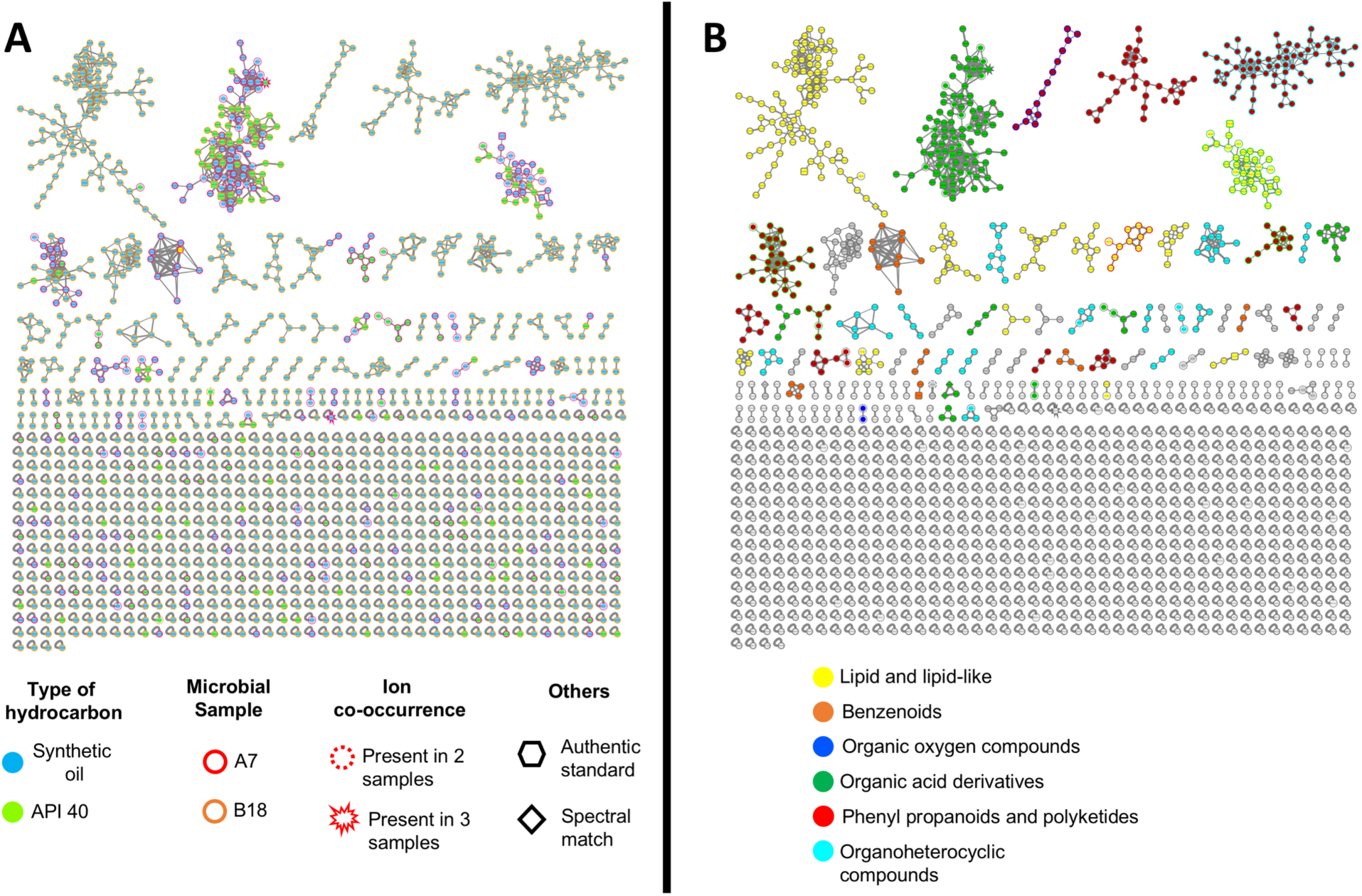
Molecular network of detected chemistries of hydrocarbon-degrading microbes from deep-marine sediments. (A) Global mass spectral molecular network of A7 and B18 grown with synthetic petroleum (SO) and petroleum API 40 (API 40) for 28 days. Each node represents a precursor ion (MS1) and edge thickness between nodes indicates similarity in MS2 fragmentation patterns. Node color, border color, and border shape indicate the type of energy used, microbial sample origin and ion co-occurrence in multiple samples, respectively. In addition, the geometrical figure (rhombus and hexagon) of the node indicates the use of an authentic standard or a spectral match (within GNPS spectral libraries). A total of 1 501 MS1 are shown in the network. (B) Structural annotation of the global molecular network of A7 and B18 at the superclass level. For each network with ≥2 nodes, a consensus candidate structure per node was assigned by NAP and each structure was subsequently classified using Classyfire, whereby the most frequent consensus classifications per network or cluster were retrieved to assign a putative superclass annotation to each network or cluster. Clusters with gray nodes indicate unassigned superclass.

**Figure 7.**
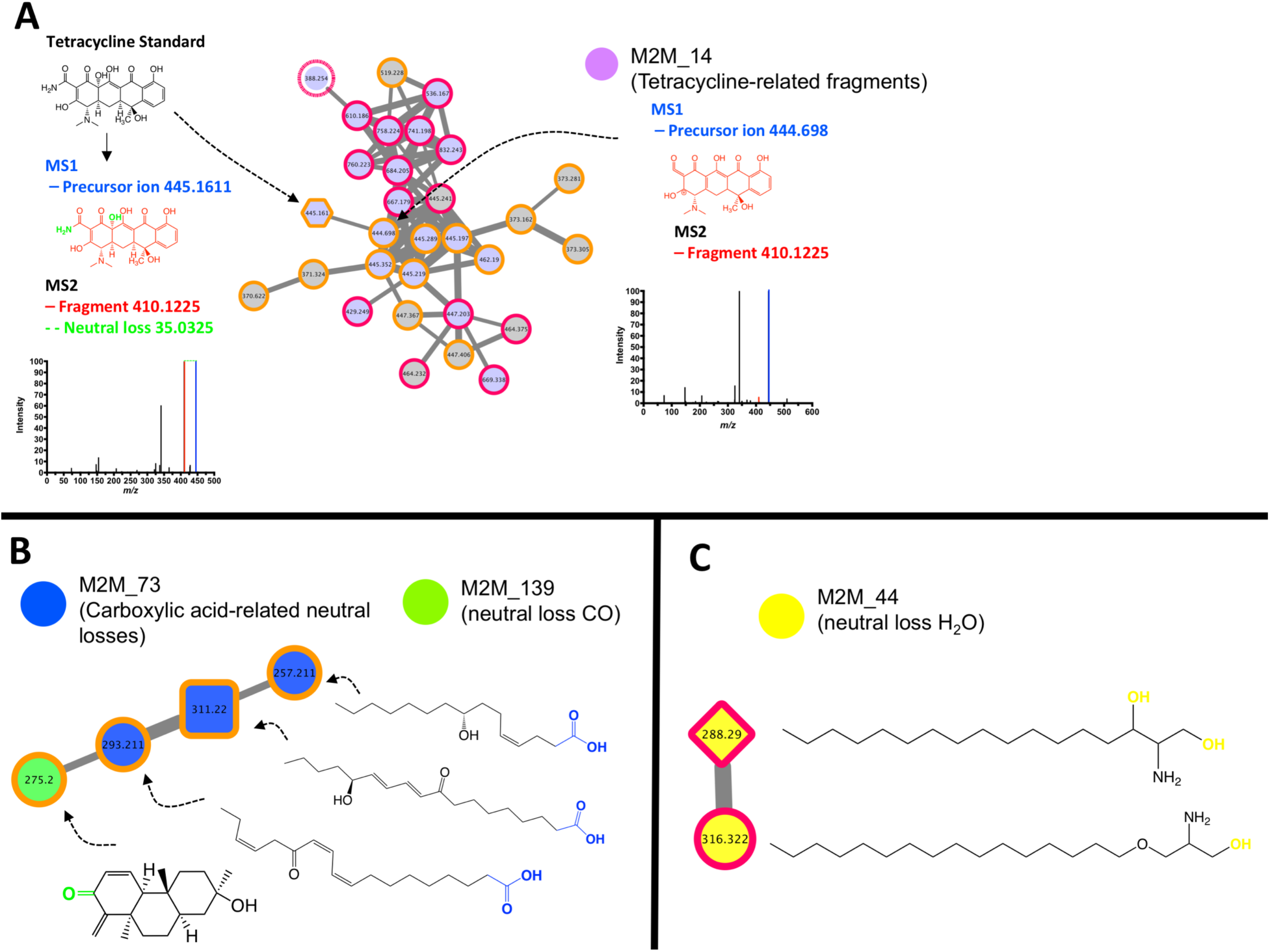
NAP/MS2LDA/MZcloud-driven metabolite annotation of selected chemistries of hydrocarbon-degrading microbes from deep-marine sediments. Mass2motifs (M2M) suggest the presence of hydrocarbon-related and non-related chemistries. (A) Tetracycline-related M2M annotated by using an authentic standard (used to enrich the global network). Fragment 410.1225 *m/z* (M2M_14) was assigned to a tetracycline-core (red substructure) using Heuristic fragmentation prediction by MZcloud (www.mzcloud.org). (B) Cluster containing the M2M_73 linked to carboxylic acids. Chemical structures drawn here are the top-ranked consensus candidates predicted by NAP. (C) Cluster containing the M2M_44 linked to neutral water loss and amine-containing fragments. Chemical structures drawn here are the top-ranked consensus candidates predicted by NAP.

MS2LDA also predicted M2M containing neutral losses of H_2_O and CHOOH hinting at the presence of alcohol and/or carboxylic acid chemistries. For instance, **Figure 7B** shows a network of 4 nodes, whereby 3 nodes are grouped within the M2M_73 containing neutral losses characteristic of carboxylic acids (i.e., loss of 18.0075 [H_2_O] and 46.0025 [CHOOH]) and the top-ranked NAP consensus structures annotates such nodes as carboxylic acids. The remaining node in such network is grouped into another M2M that contains a neutral loss of a carbonyl group and the top-ranked NAP consensus structure annotates such node as a cyclic ketone. Likewise, **Figure 7C** shows a two-node network, containing a putative annotation to C17-sphinganine (as per spectral match), grouped within the M2M_44 which contains H_2_O neutral loss and amine-containing fragments in agreement with the top-ranked NAP annotation as amine-alcohol chemistries (sphinganines).

Globally, MS2LDA analysis showed the co-occurrence of neutral losses of H_2_O, double H_2_O, NH_2_, and CHOOH suggesting the capacity of A7 and B18 to metabolize hydrocarbons into carboxylic acids and amine derivatives or amides (**Supplementary Figure 2**). For instance, the saturated dicarboxylic acids azelaic (189.1120 *m/z* [M+H]^+^) and sebacic acid (203.1284 *m/z* [M+H]^+^), associated to alkane degradation (Neil M. Broadway 1993), were detected and its annotation was confirmed by authentic standards (**Supplementary Table 1**).

## Discussion

The deep-marine biosphere, a space comprising more than 1 000 m below the sea surface, remains largely unexplored when compared to terrestrial ecosystems. Most of the microbiological and petroleum related studies, have focused on the impact of oil spills on microbial communities or on the hydrocarbon-degrading potential of microbes, primarily at the sea surface or shallow waters level (Brooijmans et al 2009). Limited studies have employed metagenomic tools to characterize deep-marine microbes and, in the best case, few have utilized hyphenated techniques (e.g., gas chromatography coupled to mass spectrometry) to evaluate their capacity to degrade hydrocarbons (Bacosa et al 2018, Kimes et al 2013). In addition, scarce information is available on the description of the full metabolic machinery of marine microbes with the potential to produce secondary metabolites when using hydrocarbons as their sole source of energy. Marine microbes have emerged as a rich biological resource of new classes of therapeutics (Williams 2009) and the analysis of genome sequence data suggest that even well-studied bacteria could produce many secondary metabolites than those discovered thus far (Cimermancic et al 2014, Doroghazi et al 2014).

The main goal of this study is to profile the metabolome derived from deep-marine microbes of the GM grown with hydrocarbons, in order to expand our knowledge about the metabolic machinery capabilities of such deep-biosphere inhabitants. Two marine samples were selected, although this will be expanded towards the mapping of the southwestern/eastern GM.

The metagenomic sequencing analysis of deep-marine sediments showed a predominant abundance of *Proteobacteria* as shown by other studies performed in GM-collected samples (Godoy-Lozano et al 2018, Mason et al 2014, Sanchez-Soto Jimenez et al 2018, Will A. Overholt 2018). Although, both microbial samples (A7 and B18) were taken from different locations, the most predominant genera (e.g., *Nitrospira*) were similar among them, but with differences in the abundance. Noteworthy, Overholt et al., suggested a consistent seafloor microbiome, predominantly linked to *Proteobacteria*, across regions of the northern and southern GM after the analysis of >700 marine sediment samples (water depth range 16-2 293 m) collected from 29 sites (Will A. Overholt 2018). Moreover, in agreement with our data, Overholt et al., demonstrated that *Nitrospira* genus was only present in deep-sediments. Noteworthy, changes in the microbial community has been noted in hydrocarbon-contaminated enviroments, for instance, a higher abundance of *Proteobacteria* has been reported in sediments exposed to the oil spill by the Deepwater Horizon disaster (Mason et al 2014). Consistently, we have noted the same pattern when environmental sediments were incubated with hydrocarbons (API 40 petroleum). The ODB associated with the *Pseudomonas* and other genera increased in abundance after 28 days of growth, as expected. It is worth mentioning that although A7 and B18 microbes grew (as per the oxygen consumption analysis) after such period using hydrocarbons as energy, a lower number of genera were detected compared to their wild initial conditions. This suggests that not all microbes are capable of growing under our *in vitro* conditions and those that survive are associated with hydrocarbon-degrading properties, including primarily *Pseudomonas, Staphylococcus, and Bacillus (Xu et al 2018).*

A considerable number of studies has documented the capacity of marine microbes to degrade hydrocarbons by inferring their degradation potential as per the presence of certain bacterial communities (i.e., ODB) (Brooijmans et al 2009, Mason et al 2014). Other studies have employed hyphenated techniques (e.g., gas chromatography/LC-coupled to MS) to study the metabolism of selective groups of hydrocarbons (Bacosa et al 2018, Kimes et al 2013), but the general knowledge of marine microbial metabolism is still limited.

Marine microbes are capable of synthesizing novel molecules, distinct to those of their terrestrial counterpart, some of them with prominent therapeutic activities (Blunt et al 2018), and are therefore, considered a relatively new and rich source of bioactive chemistries based on their metabolic machinery. It is estimated that approximately 28 500 marine natural products had been identified up to 2016 (Blunt et al 2018). However, the carbon source commonly used to feed the microbes in those studies are carbohydrates, amino acids, lipids, among other easily metabolized nutrients. To our knowledge, there is no single study evaluating the metabolome of deep-marine or shallow microbes when grown with hydrocarbons. Our study highlights, for the first time, the diversity of chemistries produced by deep-marine microbes when grown with hydrocarbons. This was achieved using an untargeted metabolomic analysis that allows collecting spectral data (MS1 and MS2) in an unbiased manner (Schrimpe-Rutledge et al 2016). We employed LC-MS/MS, which possesses certain advantages over the most commonly used technique, gas chromatography coupled to MS (GC-MS). Some of the advantages of LC-MS over GC-MS are very well described and include the capability of LC-MS to detect both volatile and non-volatile and low and high molecular weight compounds (Schrimpe-Rutledge et al 2016). In addition, complementary ‘‘state of the art’’ softwares for the analysis of mass spectral data were utilized (da Silva et al 2018, Mohimani et al 2018, van der Hooft et al 2016). The global description of the metabolome of marine microbes shows molecules with molecular masses between 114 and 883 Da with low metabolite co-occurrence or overlapping between both microbial samples and a relative low spectral match (<0.5%) against public spectral libraries (level 2 or 3 annotations according to the 2007 metabolomics standards initiative (Sumner et al 2007)). This could be attributed, in part, to the performed combined analysis of the metabolomes derived from two different carbon sources and microbial consortia (with different microbial communities). However, this approach allowed us to comprehend their metabolic machinery potential and to provide a global characterization of their metabolomes. The low spectral match we observed could be attributed to differences in the type of instrument and data acquisition parameters employed. Nonetheless, we included authentic standards (analyzed under the same conditions as our samples) with the aim to enrich the molecular network, and these when analyzed showed a correct spectral match, thereby suggesting that the relative low annotation noted is related to the absence of such metabolites within public spectral libraries. To overcome this limitation, we applied *in silico* structure prediction (i.e., NAP) to obtain *in silico* fragmentation-based metabolite annotation as reported by others (da Silva et al 2018, Kang et al 2018, Kang et al 2019). This type of analysis allows us to annotate (by a consensus candidate structure) most of the nodes (or metabolites) within the global molecular network and to assign a chemical ontology to each annotated metabolite as per ClassyFire (Djoumbou Feunang et al 2016). The chemical classification by ClassyFire consists of up to 11 different levels from a very general to a more specific classification standpoint. The most abundant superclass within each cluster represented the entire cluster in our data. Our analysis suggests a diverse type of chemistries as per the various superclasses found. The most abundant superclass was lipid and lipid-like which includes typical hydrocarbon derivatives (e.g., linear carboxylic acids, alcohols, etc.,) as those reported as petroleum products by microbial metabolism (Brooijmans et al 2009). Moreover, by using MS2LDA combined with MZcloud as a complementary tool, we were able to predict with confidence that most of the metabolites generated by A7 and B18 are carboxylic acids and alcohols. Indeed, we confirmed, by using authentic standards (level 1 annotation according to the 2007 metabolomics standards initiative (Sumner et al 2007)), the presence of azelaic and sebacic acid, two linear dicarboxylic acids reported as metabolites of the microbial degradation of alkanes (Neil M. Broadway 1993).

Due to the low level of annotation by spectral match noted herein and the complexity of microbial metabolism, selected clusters of the global network were analyzed to provide a more detailed characterization of the chemistries associated to A7 and B18 metabolism. Focus was made on 3 relevant clusters either because it contained a spectral match or an authentic standard as a node within the cluster. Likewise, as per MS2LDA and MZcloud analysis, we putatively annotated a cluster as tetracycline-related chemistries (Figure 7A). It is important to point out that we did not detect tetracycline as a metabolite, but we suggest related chemistries in such cluster due to the use of a tetracycline standard (that clustered within the network) and manual inspection of the node’s fragmentation pattern. For instance, the fragment 410.1225 *m/z* in the node connected to tetracycline standard correlates with the antibiotic scaffold predicted by MZcloud Heuristic fragmentation model. The presence of antibiotic-related metabolites in A7 and B18 microbial consortia may hint at the biological competition or antagonism exerted between diverse co-existing microorganisms (Marmann et al 2014). Noteworthy, the well-known antibiotic producer genus *Streptomyces (Watve et al 2001)* was detected in A7 and B18 samples, thereby correlating its presence to the tetracycline-related chemistries. On the other hand, the presence of sphingolipid-precursor metabolites (Figure 7B) was correlated to the presence of the genus *Sphingomonas* (Figure 5) (Harrison et al 2018). These two examples highlight the capabilities of integrating untargeted metabolomics and metagenomics to the characterization of microbes in marine consortiums.

In conclusion, this study represents the first application of molecular networking and *in-silico* dereplication to the metabolome associated with the degradation of hydrocarbons by microbes obtained from the deep-marine biosphere. The use of complementary *in silico* dereplication strategies such as NAP and MS2LDA is capable of providing a global perspective of the metabolic potential of hydrocarbon-degrading microbes by inventorying the chemistries generated from a general to a specific standpoint. Our study evidences the capabilities of marine microbes to synthesize diverse chemistries when hydrocarbons are used as the sole source of energy, which may hold a role for the discovery of new bioactive chemistries.

## Acknowledgements

We thank the crew of the GM oceanographic campaign XIXIMI-4 who participated in sample collection.

