## Supplementary Information for "Metabolic and metagenomic profiling of hydrocarbon-degrading microorganisms obtained from the deep biosphere of the Gulf of México"

***for***

**Supplementary Table 1. Authentic standards used for metabolite characterization or molecular network enrichment**

| Compound | Exact mass  [M+H]+ | Experimental  [M+H]+ | Mass accuracy (ppm) | Sample where detected |
| --- | --- | --- | --- | --- |
| Azelaic acid | 189.1127 | 189.1120 | -3.7 | B18-API40 (NH_4_^+^ adduct), A7-PS, B18-PS |
| Sebacic acid | 203.1284 | 203.1280 | -1.9 | B18-API 40, A7-PS, B18-PS |
| Tetracycline | 445.1611 | 445.1610 | -0.2 | Standard used for network enrichment |
| Amoxicilin | 366.1124 | 366.1120 | -1.09 | Standard used for network enrichment |
| Levofloxacin | 362.1516 | 362.1510 | -1.6 | Standard used for network enrichment |
| Ceftriaxone | 555.0539 | 555.0540 | 0.18 | Standard used for network enrichment |

**Supplementary Figure 1.** Molecular network of A7 and B18 microbial consortia grown with hydrocarbons and stratified by the mass-to-charge ratio of the precursor ion. The size of the node denotes the mass-to-charge of the precursor ion.

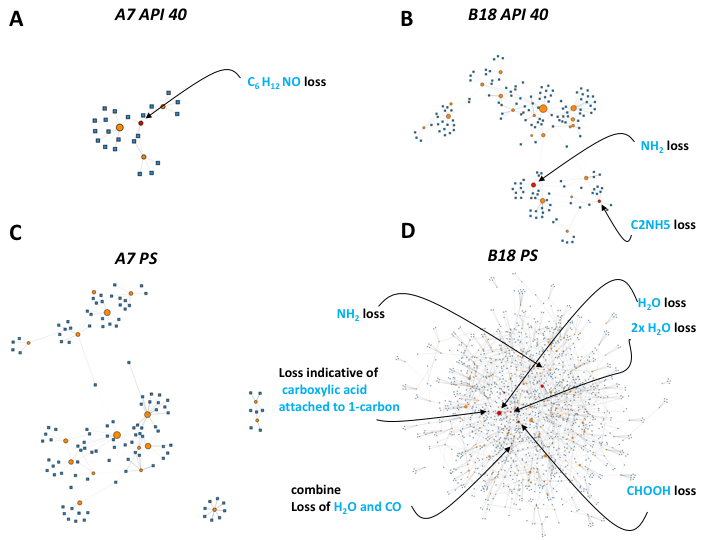

**Supplementary Figure 2.** Network visualization of the Mass2Motifs (M2M) associated to individual marine microbial metabolomes. M2M containing at least 5 molecules are shown for A7 (A and C) and B18 (B and D) consortia grown with API 40 or synthetic petroleum (PS). Nodes represent M2M (circle) and molecules (squares). Red nodes indicate a M2M match against the MassBank M2M database.
